# Nested contextual change and the temporal compression of episodic memory

**DOI:** 10.64898/2026.02.26.708184

**Authors:** Matthew Logie, Camille Grasso, Virginie van Wassenhove

## Abstract

How does the structure of events influence the when and the where of life experiences in comparison to the what? We developed a novel virtual reality (VR) environment to understand how the quantity of information within nested structures influence participants’ memory for events. Participants moved through a series of virtual rooms (events) where images (items) appeared in randomised locations on a 3 by 3 grid located on a wall. Participants were asked to remember the what (old/new), when (timeline location), and where (grid location), of the images they experienced. Two types of nested events were tested (6 rooms, each containing 4 images; 3 rooms, each containing 8 images), all lasting an equivalent number of seconds. We found a strong temporal compression effect at nested levels in which participants remembered early items and events happening later, and later items and events happening earlier, than the original experience. Crucially, presenting four-item events resulted in a greater compression rate than eight-item events. We also found greater temporal distances between pairs of items occurring within eight-item events than pairs of items which occurred on either side of a boundary. Memory for when depends on the nested compression of information within events.

## 1. Introduction

Our experience of time fluctuates based on our perception of change ^1,2^ and may fundamentally rely on episodic memory, that is, our capacity to remember the what, when, and where of specific experiences ^3,4^. From this perspective, the subjective experience of time is not due to a direct encoding of durations, but rather through the construction, storage and reconstruction of information within events ^5,6^. Event boundaries, provided by large contextual changes, naturally serve as breakpoints that organize the “stream of consciousness” into chunks or experiential episodes. The information stored within an event can vary in terms of both accuracy and precision: for example, correctly remembering an encountered item (‘what’) but being unable to identify the ‘when*’* or the ‘where*’* of that encounter. In the current work, we ask how the structure of events influences the ‘when’ in comparison to the ‘what’ and ‘where’ of episodic memory. Specifically, we ask how the quantity of information between event boundaries influences the ability to place when an item occurred on a timeline. We expect to find variations in temporal compression, response times and distances between items in memory for ‘when’ without equivalent variations in memory for ‘what’ or ‘where’.

From the episodic perspective, the encoding of information involves the variable compression ^7,8^ of events constructed within working memory ^9^. Working memory can both construct new events and reconstruct past events with varying levels of accuracy and precision ^10,11^. These varying levels of compression are expected to govern the temporal distances between items stored in episodic memory. Information is constructed within working memory and experiencing an event boundary, defined as a prediction error or as an accumulation of change reaching a threshold ^12–14^, drives the compression and encoding of information into episodic memory. In turn, information encoded into long-term memory is only accessible through the reconstruction of an event within working memory ^9,15,16^.

The scanning time (aka response times) between events in memory have been shown to depend on the number of event boundaries ^17^. In this study, participants viewed a short movie. Afterwards, a scene was described (e.g. “remember when you saw flames in the hallway”) and participants were asked to describe the next scene containing a particular event (e.g. “when is the next time you see fire”). The time it took a participant to answer (response time) was slower with the greater number of event boundaries between the scenes. The results were explained by a skipping process that quickly ‘skips’ to an event boundary followed by a slow scanning of memory to identify a particular item within an event, starting from the first item. This constitutes a prediction for our study: the quantity of information within events should influence the response times when participants are asked to place items on a timeline. More specifically, if participants scan memory from the beginning of an event, response times for the first item within an event should be faster than items occurring at the end of an event.

Temporal compression has previously been demonstrated in terms of a shorter response time to replay episodes from memory than the original experience. While previous work has demonstrated temporal compression effects at a single level based on increasing the number of seconds of presentation with movies ^18^, here, we focus on the quantity of information within nested events while maintaining the same total number of seconds for the overall sequence. If temporal compression depends on the quantity of information, then adjusting the structure of nested events will result in differing compression effects even when there is no difference in the total number of seconds. Adjusting how many events occur within the same number of seconds also allows for the evaluation of temporal compression in terms of errors of temporal distances between items. We predict that errors of temporal displacements and response times will depend not only on distance (in terms of number of items) to the closest event boundary, but also on the distance to either the very beginning or the end of a presented sequence. Additionally, the repetition of events can lead to events becoming more predictable, allowing for more efficient compression for later events than early events. Increasing the repetition of events may also lead to a shortening of the passage of time ^31^.

Episodic memory also depends on nested structures characterised by different timescales ^19^. Previous studies of event segmentation have identified a nested structure of events occurring at fine and coarse grains ^6,20,21^. For instance, experiencing a context shift, such as travelling through doorways ^22^ to a new virtual location provides an event boundary that defines the end of an event ^23^. Similarly, turning corners while moving through a real or virtual environment provides boundaries, and the number of turns experienced influences rates of compression ^24^: an expansion of time occurs when experiencing contextual changes provided by turning corners ^25^ or increasing the number of events in virtual environments ^26^. Duration estimates expand when experiencing unpredictable events ^26–28^, thus temporal distances in memory for ‘when’ should also depend on the structure of events.

In addition to a segmentation account, contextual changes defined in models of contextual drift provide an explanation of episodic memory performance ^29^. One account of contextual drift defines an event in terms of a period of stable context drift occurring between accelerated context shifts ^30^ and another proposes a partial resetting and reinstatement of context representations at event boundaries ^31^. Accelerating rates of change at boundaries (such as the transitions between virtual rooms, which we will be exploring here) increase the temporal distance between events in terms of how much contextual change has occurred. While previous work has focused on explaining memory for ‘what’ and the temporal order of items within events ^30,31^, models of context drift also inform predictions for how boundaries affect subjective temporal distances between items.

The model of ‘resetting’ context ^31^ defines temporal distances as the dissimilarity in context between target items and a start of sequence context (i.e. item 1 in event 1). A partial reinstatement of context at boundaries brings the start of each subsequent event closer in similarity and therefore would predict smaller temporal distances for pairs of items occurring across events than for pairs of items occurring within events. Whereas for the model describing an acceleration of context drift at boundaries, temporal distances are determined by a direct comparison of contextual similarity between target items ^30^. An acceleration at boundaries reduces the similarity of context for pairs occurring across events and would predict larger temporal distances for across event item pairs than within event item pairs. Note that both variations of the temporal context model predict equivalent distance between items within events as seen in Figure 1.

**Figure 1:**
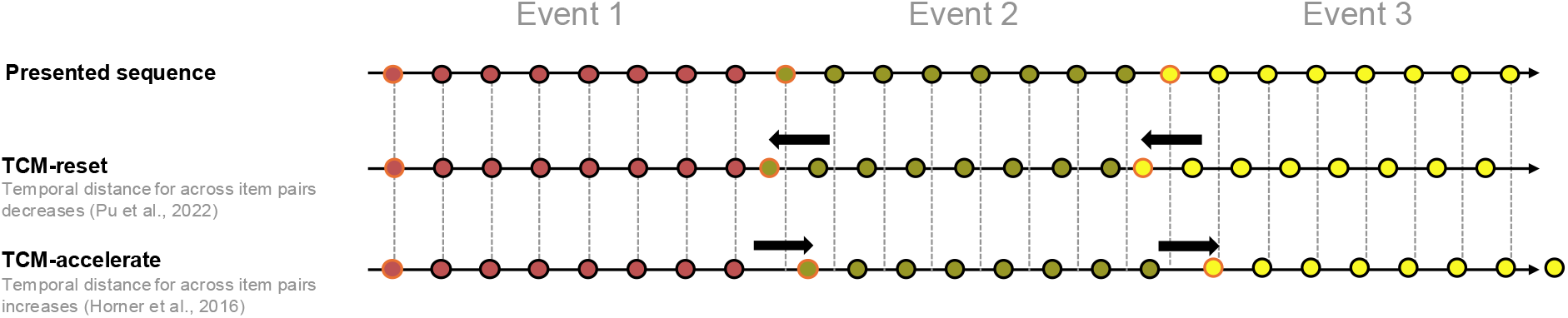
Predictions for temporal distances based on event structure. TCM-reset (top): Event boundaries (red highlights) pull events closer together (arrows). TCM-acceleration (bottom): Event boundaries push events further apart.

Altogether, we predict that the quantity of information within events will determine the compression rate and drive the accuracy and similarity of temporal displacement errors. To test this prediction, we exploited movement transitions as boundaries within a novel virtual reality (VR) environment to provide nested event structures containing different quantities of information (number of images). Our main theoretical question is to ask how the quantity of information within events influences memory for ‘when’ in comparison to memory for ‘what’ and ‘where’. Specifically, we asked whether memory for ‘when’ varies based on the quantity of information within events and the compression and reconstruction of information, even when there are no differences in the total number of seconds that pass. To answer this question, we compared temporal responses, temporal displacement errors, spatial displacement errors, recognition performance and associated response times based on the position of to-be remembered images (items) within a segmented sequence of virtual rooms (events).

## 2. Methods

### 2.1. Participants

The sample size was chosen based on previous work identifying an influence of event boundaries on temporal displacement errors ^32^ and on previous work comparing memory performance for items occurring within segmented virtual rooms ^33^. Twenty-eight participants (M = 25.78 years old, SD = 4.77) took part in the study. Two participants were recruited but did not complete the task. Hence, 26 participants were included in the final analysis (M = 26.04 years old, SD = 4.85). We also conducted a power analysis for 26 subjects each completing 3 trials containing 24 images (72 images per condition) providing a total of 1872 observations per condition. The responses were averaged to produce a single value per participant per position, providing a total of 624 observations. We identified an intraclass correlation of 0.611 [CI= 0.575, 0.644] for the ‘when’ response, our primary measure of interest. The design allows for identifying small effects of f = 0.1 at over 95% power. The code used for conducting the power analysis is available at the OSF link at the bottom of the manuscript. All participants were informed about the study requirements and signed a written consent form in agreement with the Declaration of Helsinki (2013) ^34^. All experimental protocols were approved by the local Ethics Committee on Human Research (CEA-CPP-100049).

### 2.2. Materials

The VR environment was constructed with Unity and run with the HP Reverb G2 Omnicept VR headset. The VR environment consisted of a series of colored rooms (Figure 1A). Within each virtual room (event), a series of images (items) of unrelated everyday objects were displayed in a random order at different locations on a 3-by-3 grid presented on the wall (Figure 1A, right panel). The images were unique and not repeated across trials. The experiment consisted of 6 blocks each containing 24 images for a total of 144 ‘old’ images. For each participant the images were shuffled once at the start of an experiment and presented in a random order (i.e. the first group of 24 from the shuffled list were presented in the first block, the second group of 24 were presented in the second block). An additional 144 ‘new’ images were employed to test recognition memory which were also shuffled at the start of an experiment to produce a random order for each participant. The first 24 images of the ‘new’ list were combined with the first 24 images of the ‘old’ list and again shuffled to produce a randomised list of 48 images for the test phase. The combining and shuffling of old and new lists occurred again for each subsequent test phase. The design ensured that each participant saw the same images in different randomised orders. Images were taken from the Bank of Standardized Stimuli (BOSS) database ^35^.

### 2.3. Procedure

#### 2.3.1. Practice sessions

Prior to the start of the study, a practice session was provided. The practice session consisted of a single trial of the study and test phase (see 2.3.3) using only one room. Participants could move on to the study phase if old items could be recognised, otherwise another practice trial was run.

#### 2.3.2. Tolerance to Virtual Reality

The practice was followed by a simulator sickness questionnaire ^36,37^ testing participants sensitivity to VR. A score of less than 50 based on the original scoring ^37^ was required for participants to start the main experiment. No participant reached a score above 50 and all participants were thus tested.

#### 2.3.3. Experimental design

The experiment consisted of six blocks each containing two experimental phases: the study and the test phase. In each study phase (Figure 2B), the floors of each virtual room were shown in a different color (e.g. Figure 2A) with transitions between rooms providing context shifts (boundaries) between sequences of items. Each study block employed a different set of colors. Participants were automatically moved through the VR rooms, and they were instructed to remember the ‘what, when, and where’ of images presented along the sequence of virtual rooms. Two conditions were tested by block: in one condition, participants were moved through six rooms and presented with four images in each room (Figure 1A, left panel). In the other condition, participants were moved through three rooms and presented with eight images in each room (Figure 1A, middle panel). Hence, in both conditions, twenty-four images were presented by block. The full experiment consisted of a total of six blocks (three of the 6 events x 4 items, and three of the 3 events x 8 items). The order of the blocks was counterbalanced within participants. The duration of the study phase lasted the same total number of seconds (i.e. 64.8s) in all blocks and in both conditions.

**Figure 2:**
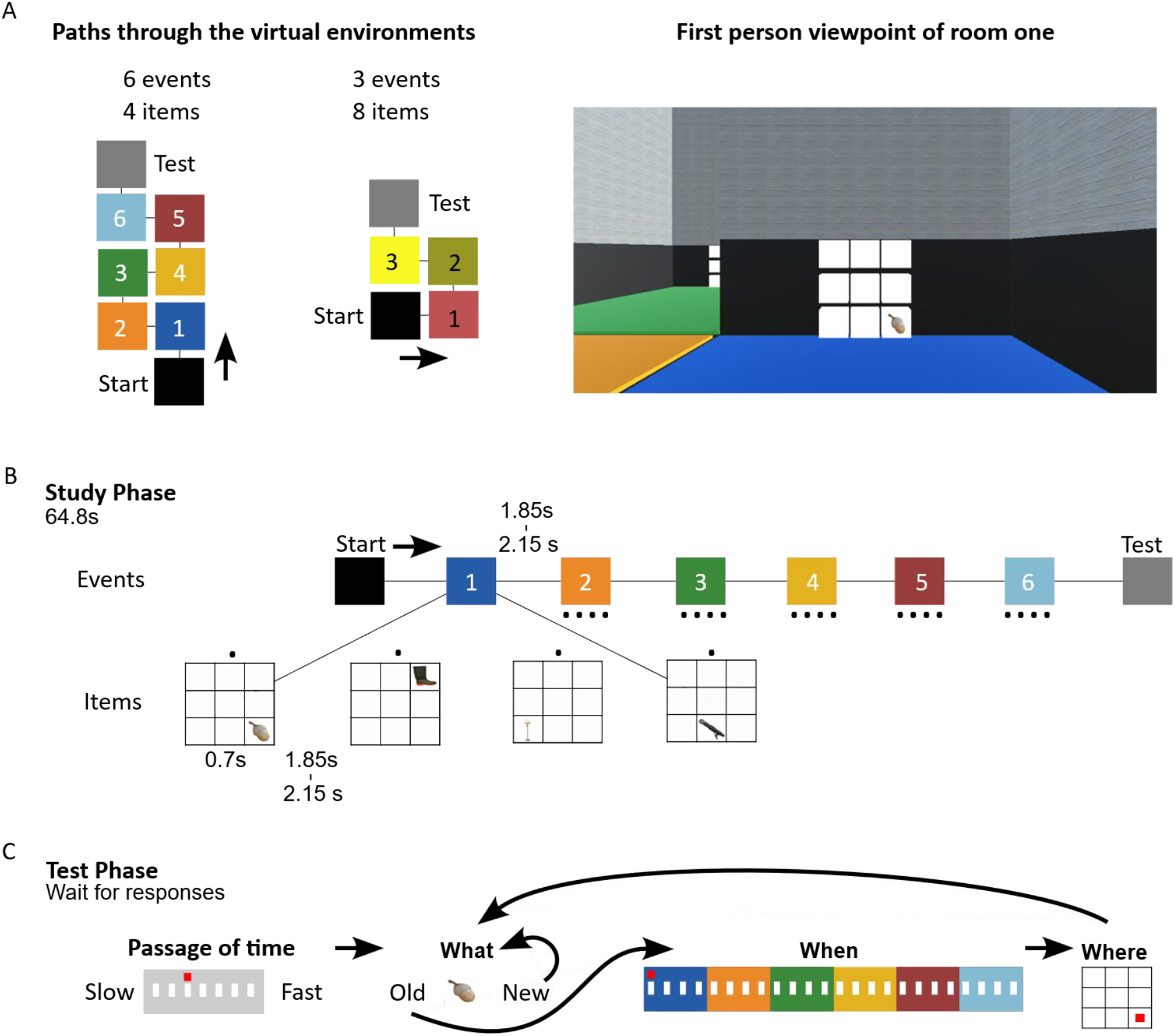
Task design. **Panel A**: Overview of task structure displaying a map of the virtual rooms or events in our design (left panel). Participants start in the black square; the arrow indicates the movement to the first room. Participants are automatically moved through rooms 1-6 (left panel) or rooms 1-3 (middle panel) until the test phase. In each room (right panel), participants see a 3-by-3 grid (white squares) on which images (items) are being presented. Half the participants completed the four-item conditions first; half the participants completed the eight-item condition first. **Panel B**: Study phase. In each room, images were sequentially presented at random locations on the grid. The study phase presented a total of 24 images (4 in each of the 6 rooms or 8 in each of the 3 rooms). Each condition took the same number of seconds to complete**. Panel C**: The test phase included a sequence of tests presented in the same order each time: a passage of time judgement, an old or new recognition test (‘what’), a judgement of ‘when’ and a judgement of ‘where’. To answer the passage of time, ‘when’ and ‘where’ tests participants used a stick on a VR controller to move a marker to the appropriate position.

After completing the study phase, participants were moved through a doorway and experienced a brief break before being asked several tests (Figure 2C). The test phase took part after each study phase for a total of 6 blocks (3 per condition) and consisted of a randomised list of 24 old images and 24 new images. First, participants were asked to make a passage of time judgement and to answer the question, “how fast did time pass since the start of the sequence?” using a Likert scale ranging from 1-7 (very slow to very fast). Following their judgement of the passage of time, participants were provided with a recognition test (‘what’ judgment), in which they were asked to judge if the image presented on the screen was ‘old’ or ‘new’. If an image was ‘old’, they were asked to perform a judgement of ‘when*’* by placing a marker at the position of the presented image on a timeline. Then, they were asked to judge ‘where*’* an image appeared and were instructed to place a marker at the location of the image on the 3x3 grid. Following the test phase, participants were then placed into the next study block and pressed a button on the VR controller to begin.

### 2.4. Analysis

In the old/new test, recognition performance was evaluated by calculating d’ prime scores, based on the proportion of previously presented items correctly recognised (hits) and the proportion of new items that were recognised as being previously presented (false alarms). Recognition performance in the ability to distinguish old and new items was evaluated with d prime. Recognition and passage of time judgements were analysed with Bayesian paired sample t-tests.

To evaluate how the nested structures influenced the performance of memory for ‘when’, ‘where’ and ‘what’ judgements based on both room (event) and image (item) position, we employed linear mixed models with likelihood ratio tests in JASP ^38^. Each linear mixed model tested for fixed effects of event position (1-6 or 1-3) and item within event position (1-4 or 1-8). The model included a random effect grouping factor of participant (1-26) with random intercepts. Note that random slopes for event and item position were not included due singularity issues in the model fit. Possible models were evaluated based on Akaike Information Criteria (AIC). The data for the ‘when, where and what’ judgements were taken from the correct recognition of an image (the hits), as not every image was correctly recognised, some data was missing from random item positions. The ‘when, where and what’ judgements were solely collected for images that were correctly recognized as ‘old’ (hits in the old/new task). Thus, images that were not recognized as old were missed and were not included in the data when quantifying the where and when of these images. The number of observations included have been specified for each analysis. The missing where and when data were randomly interspersed. To analyse differences between event and item positions, we performed post-hoc, holm corrected t-tests. Memory for ‘when’ was scored based on participants’ response position on the timeline and the temporal displacement errors. Temporal displacement errors were obtained by subtracting the expected position of an item on the timeline as presented in the study phase from the position of an item reported by the participant on the timeline in the test phase.

Memories for ‘where’ were first analysed with Bayesian one-sample t-tests against chance performance. Chance performance for placing an item on the 3 by 3 grid was 0.11 (1/9). The proportion of correct responses in memory for ‘where’ were also submitted to linear mixed models to evaluate the effect of event and item within event position on the proportion of correct placements on the grid.

Memories for ‘what’ were also analysed with linear mixed effects models based on the proportion of hits in the recognition test. For each judgement of ‘when’, ‘where’ and ‘what’ the associated response times were submitted to linear mixed models with fixed effects of event and item within event position. Response times greater than 2.5 standard deviations above the mean were excluded from the response time analysis ^17^ and the remaining response times were log transformed and z-scored.

To evaluate how the event structures influenced temporal distances between items in memory for ‘when’, temporal distances were calculated between pairs of items within or across events in each condition. Distances were calculated by subtracting the response position of the first item from the response position of the second item for each pair. The approach meant that negative distances indicate an incorrect temporal order with the second item placed earlier than the first item within the pair (incorrect distance and order). For the four-item condition, we calculated and compared distances for pairs of items within events to pairs of items across events. Each pair was separated by 1 item. For the eight-item condition we calculated distances for pairs occurring early within an event, late within an event and across an event. Each pair was separated by 3 items. The distances for each pair were analysed based on Bayesian paired sample t-tests in JASP ^38^. Evidence for either an alternative or the null hypothesis was obtained based on a default Cauchy prior with width of 0.707 ^39^. Bayes factors less than 0.33 indicate moderate evidence in favour of the null while Bayes factors greater than 3 indicate moderate evidence in favour of the alternative.

## 3. Results

### 3.1. Memory for ‘when’. Do nested event structures influence temporal responses, displacement errors and response times?

We questioned how the different event structures would influence memory for ‘when’ based on the position of an item within an event (image in the room) and the position of an event within a study block (the position of a room in the full sequence of VR rooms). To examine the role of event structure on memory for ‘when’, we employed linear mixed models (see Analysis). The analysis revealed extreme evidence for temporal compression effects (both for images within rooms and rooms within study blocks), as illustrated in Figure 3. Images occurring later in a sequence were placed earlier on the timeline. Response times were slower for first rooms than later rooms. We describe these effects below.

**Figure 3:**
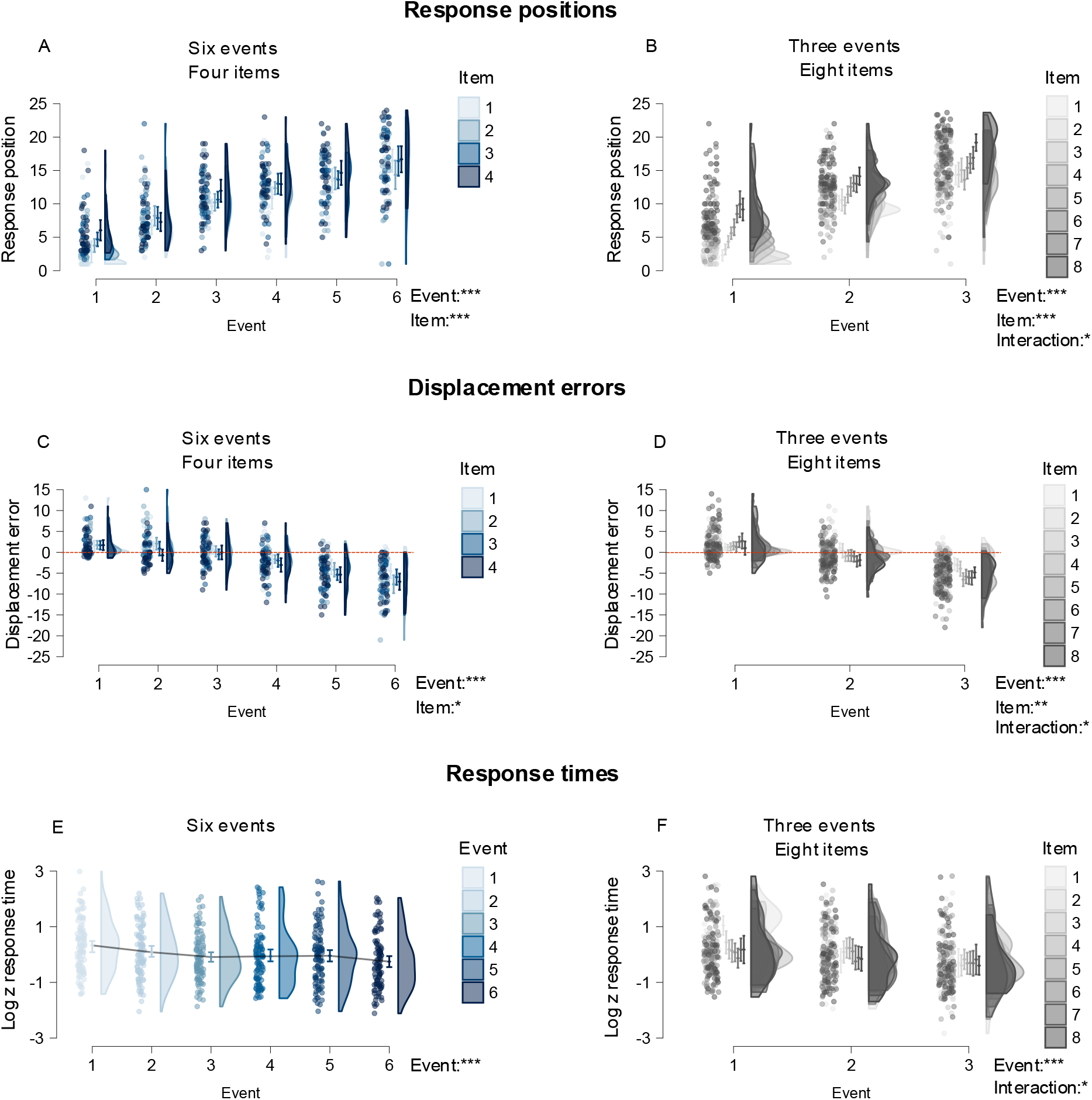
Response positions, Temporal displacement errors and Response times for judgements of ‘when’. **Panel A, Panel B**: Raincloud plots of response positions for correctly recognised images for four items per event in six events (**A**) and eight items per event in three events (**B**). **Panel C, Panel D:** Raincloud plots of temporal displacement errors for correctly recognised images for four items per event in six events (**C**) and eight items per event in three events (**D**). The dotted red lines indicate the positions for a correct response. **Panel E, Panel F**: Log transformed, z-scored response times in judgments of ‘when’ for four items per event in six events (**D**) and eight items per event in three events (**F**). Plots display overestimations of early items and underestimations of late items for temporal positions and displacement errors in both conditions at the level of both item within an event and an event within a study block (**A, B, C, D**). Response times for later events were faster than response times for early events (**D, F**) Errors in memory for ‘when’ defined as the number of items between the position of presentation and the position of response. Dots represent averaged responses from 3 blocks per participant. Error bars represent 95% confidence intervals.

#### 3.1.1. Temporal compression of memory for ‘when’

To evaluate how the quantity of information within the nested event structures influenced the accuracy of temporal responses, linear mixed models were fitted with dependent variables of response positions for both the four-item and the eight-item conditions. The model included fixed effects of event (1-6 for the four-item condition and 1-3 for the eight-item condition) and item position (1-4 for the four-item position and 1-8 for the eight-item condition).

For the four-item condition, the analysis included 597 observations (27 missing) from 26 participants (Figure 3A). The model including fixed effects of both event and item position (log likelihood = −1628, AIC = 3277) outperformed a model containing a fixed effect of event position alone (log likelihood = −1636, AIC = 3288) and a model containing an interaction between event and item position (log likelihood = −1620, AIC = 3291). The analysis revealed extremely strong evidence for a main effect of both event position, (χ^2^(5) = 452.23, p < .001) and item position (χ^2^(3) = 16.96, p < .001) on the temporal response positions. For the fixed effect estimates, the grand mean intercept across all events and items was (b = 10.77, SE = 0.17, t (25.58) = 62.48, p < .001). The post-hoc holm-corrected t-tests between first and last events showed an earlier timeline response for Event 1 (b = 4.33, CI = 3.60, 5.05) than Event 6 (b =16.00, CI = 15.25, 16.74), with an estimated mean difference of 11.67 (z = 22.55, p < .001, CI = 10.66, 12.68). The post-hoc t-tests between first and last items also showed an earlier response for Item 1 (b =9.95, CI =9.34, 10.56) than Item 4 (b = 11.617, CI = 11.01, 12.23) with an estimated mean difference of 1.67 (z = 3.93, p < .001, 0.84, 2.50). The results show that participants could place the events and the items within events in the correct temporal order, with the last events and the last items placed later than the first events and the first items, respectively. The response positions were compressed towards the midpoint of the sequence, consistent with central tendency effects in timing ^40^.

For the eight-item condition, the analysis included 619 observations from 26 participants (Figure 3B). The model containing an interaction between event and item position (log likelihood = −1614, AIC = 3280) outperformed a model containing fixed effects of event and item position (log likelihood = −1633), AIC = 3291) and a model including only event position (log likelihood = −1714, AIC = 3439). The analysis revealed extremely strong evidence in favour of a main effect for both event position, (χ^2^(2) = 543.5, p < .001), item position (χ^2^(7) = 171.12, p < .001) and an interaction between event and item position (χ^2^(14) = 38.74, p < .001) on the temporal response positions. For the fixed effect estimates, the grand mean intercept across all events and items was (b = 11.21, SE = 0.20, t (25.92) = 56.69, p < .001). The post-hoc holm-corrected t-tests between first and last events showed an earlier response for Event 1 (b = 6.19, CI = 5.66, 6.72) than Event 3 (b = 15.61, CI = 15.08, 16.14) with an estimated mean difference of 9.42 (z = 29.64, p < .001, CI = 8.80, 10.04). The post-hoc t-tests between first and last items also showed an earlier response for Item 1 (b = 8.94, CI =8.17, 9.71) than Item 8 (b = 14.16, CI = 13.38, 14.93) with an estimated mean difference of 5.22 (z = 10.11, p < .001, CI = 4.21, 6.23). The effect of item positions within events on the reported position of items on the timeline produced a much larger effect in the eight-item condition than in the four-item condition.

Altogether, we found that the temporal positioning of items on the timeline reported by participants showed a temporal compression effect. This temporal compression effect occurred at nested levels both for events and items within events. Strikingly, we found an interaction between event and item position when presenting eight items per event in three events but not when presenting four items per event in six events.

#### 3.1.2. Error displacements reflect nested temporal compression

Next, we questioned the influence of nested event structures on temporal displacement errors. In particular, we asked whether participants showed less errors for items that occurred closer to event boundaries ^41^. The temporal displacement errors were scored by subtracting the presented position of an image in the study phase from the response position of an item in the test phase. For example, an image presented in position 4 in the study phase and placed on the timeline in position 2 in the test phase would be scored as −2. The analysis of temporal displacement errors followed the same approach as for the response positions: separate linear mixed models were fitted with temporal displacement errors as dependent variables for the four-item and the eight-item condition.

For the four-item condition, the analysis included 595 observations (Figure 3C). The model including fixed effects of both event and item position (log likelihood = −1602, AIC = 3227) outperformed a model containing fixed effects of event position alone (log likelihood = −1607, AIC = 3231) and a model containing an interaction between event and item position (log likelihood = –1594, AIC = 3239). The analysis revealed extremely strong evidence in favor of an effect of both event position, (χ^2^(5) = 303.86, p < .001), and item position (χ^2^(3) = 10.14, p = .017) on temporal displacement errors. For the fixed effect estimates, the grand mean intercept across all events and items was (b = −1.74, SE = 0.17, t (25.74) = −10.38, p < .001). The post-hoc holm-corrected t-tests between first and last events showed overestimated displacement errors for Event 1 (b = 1.67, CI = 0.97, 2.38) and underestimated errors for Event 6 (b = −6.27, CI = −7.0, −5.54), with an estimated mean difference of −7.94 (z = −15.78, p < .001, CI = −8.93, −6.96). The post-hoc t-tests between first and last items showed less underestimation of errors for Item 1 (b = −1.12, CI = −1.71, −0.53) than Item 4 (b = −2.39, CI = −2.98, −1.80) with an estimated mean difference of −1.27 (z = −3.09, p = .002, CI = −2.08, −0.47).

Hence, both event position and item positions within events had an extremely large influence on participants’ temporal displacement errors. These results explicitly identified the first event being placed later on the timeline, and the last events being placed earlier on the timeline than would be expected based on the original presentation.

For the eight-item condition, the analysis included 612 observations (Figure 3D). The model including an interaction between event and item position (log likelihood = −1592, AIC = 3235) outperformed a model containing fixed effects of event and item position (log likelihood = −1611, AIC = 3246) outperformed a model containing only event position (log likelihood = −1621, AIC = 3252). The analysis revealed extremely strong evidence for an effect of both event position (χ^2^(2) = 321.79, p < .001) and strong evidence for item position (χ^2^(7) = 22.77, p = .002) and an interaction between event and item position (χ^2^(14) = 38.54, p = .001) on temporal displacement errors. For the fixed effect estimates, the grand mean intercept across all events and items was (b = −1.37, SE = 0.20, t (25.97) = −6.73, p < .001). The post-hoc holm-corrected t-tests between first and last events again showed overestimated displacement errors for Event 1 (b = 1.59, CI = 1.06, 2.12) and underestimated displacement errors for Event 3 (b = −4.90, CI = −5.44, −4.36) with an estimated mean difference of −6.49 (z = −20.48, p < .001, CI = −7.10, −5.87). The post-hoc t-tests between first and last items also showed less underestimation of errors for Item 1 (b = −0.18, CI = −0.96, 0.60) than Item 8 (b = −1.90, CI = −2.67, −1.13) with an estimated mean difference of −1.72 (z = −3.34, p = .001, CI = −2.73, −0.71).

Thus, the eight-item condition also showed significantly large effects of event and item positions on the temporal displacement errors both for events and items within events. While no interactions were found when presenting four items per event in six events. The analysis of displacement errors also revealed an interaction between event and item position when presenting eight items per event in three events. The errors in memory for when depend on the quantity of information within nested event structures.

#### 3.1.3. Response times support the backward scanning of memory for ‘when’

To examine the role of event structure on memory scan times in memory for ‘when’ we analysed the response times for correctly recognised images.

For the four-item condition, the analysis included 595 observations (Figure 3E). The model including fixed effects of event position alone (log likelihood = −724.6, AIC = 1465) outperformed a model with fixed effect of both event and item positions (log likelihood = −723.8, AIC = 1470) and a model containing an interaction (log likelihood = −713.3, AIC = 1497) . The analysis revealed strong evidence for an effect of event position on response times (χ^2^(5) = 28.64, p < .001). Fixed effect estimates identified a grand mean for transformed response times (b = −0.008, SE = 0.13, t (25.97) = −0.06, p = .95). The post-hoc holm-corrected t-tests showed a slower response time for Event 1 (b = 0.28, CI = 0.01, 0.56) compared to Event 6 (b = −0.25, CI = −0.54, 0.03), with an estimated mean difference of −0.54 (z = −5.00, p < .001, CI = −0.75, −0.33). The post-hoc t-tests between first and last items did not provide evidence for slower response times for Item 1 (b = 0.05, CI = −0.20, 0.29) than Item 4 (b = −0.07, CI = −0.37, 024) with an estimated mean difference of −0.11 (z = −1.05, p = .29, CI = −0.32, 0.10). Thus, response times for the first event were slower than response times for the later events. There were otherwise no differences in response times between items within events.

For the eight-item condition the analysis included 587 observations (Figure 3F). The model including an interaction between event and item position (log likelihood = −711., AIC = 1475) outperformed a model containing fixed effects of event position alone (log likelihood = −734, AIC = 1478) and a model containing fixed effects of both event and item positions (log likelihood = −729.1, AIC = 1482). The analysis revealed strong evidence for an effect of event position, (χ^2^(2) = 48.88, p < .001) and an interaction between event and item position (χ^2^(14) = 35.62, p = .001) but no effect of item position (χ^2^(7) = 9.27, p = .234) on response times. Fixed effect estimates identified a grand mean for log transformed response times (b = 0.005, SE = 0.12, t (25.84) = 0.05, p = .96) and a significant fixed effect for Event 1 (b = 0.28 (SE = 0.04, t (561.18) = 6.26, p < .001). The post-hoc holm-corrected t-tests between first and last events showed a decrease from Event 1 (b = 0.29, CI = 0.04, 053) to Event 3 (b = −0.27, CI = −0.52, −0.03) with an estimated mean difference of −0.56 (z = −7.15, p < .001, CI = −0.71, −0.40). The post-hoc t-tests between first and last items also showed a slower response times for Item 1 (b = 0.16, CI = −0.12, 0.44) than Item 8 (b = −0.12, CI = −0.40, 0.16) with an estimated mean difference of −0.28 (z = −2.16, p = .031, CI = –0.52, −0.03). Response times for the first event were slower than response times for later events. Unlike the previous condition, response times for the first item within an event were slower than the last item within an event.

Altogether, we found that response times fastened in the course of the VR sequence, with events closer to the moment of recognition being faster to recall than events earlier in the VR sequence. This pattern of results supports a backward scanning of memory^44^, with faster response times for the last item than the first item, in contrast with a forward scanning predicted by the skipping to event boundary model ^18^. The results also support a reconstruction of memory from the end of the sequence and the end of an event, with slower response times following an event boundary and faster response times before the event boundary at both event and item within event levels. Consistent with the analysis of response positions and displacement errors we found an interaction between event and item position for response times when presenting eight items per event in three events.

**Table 1.**
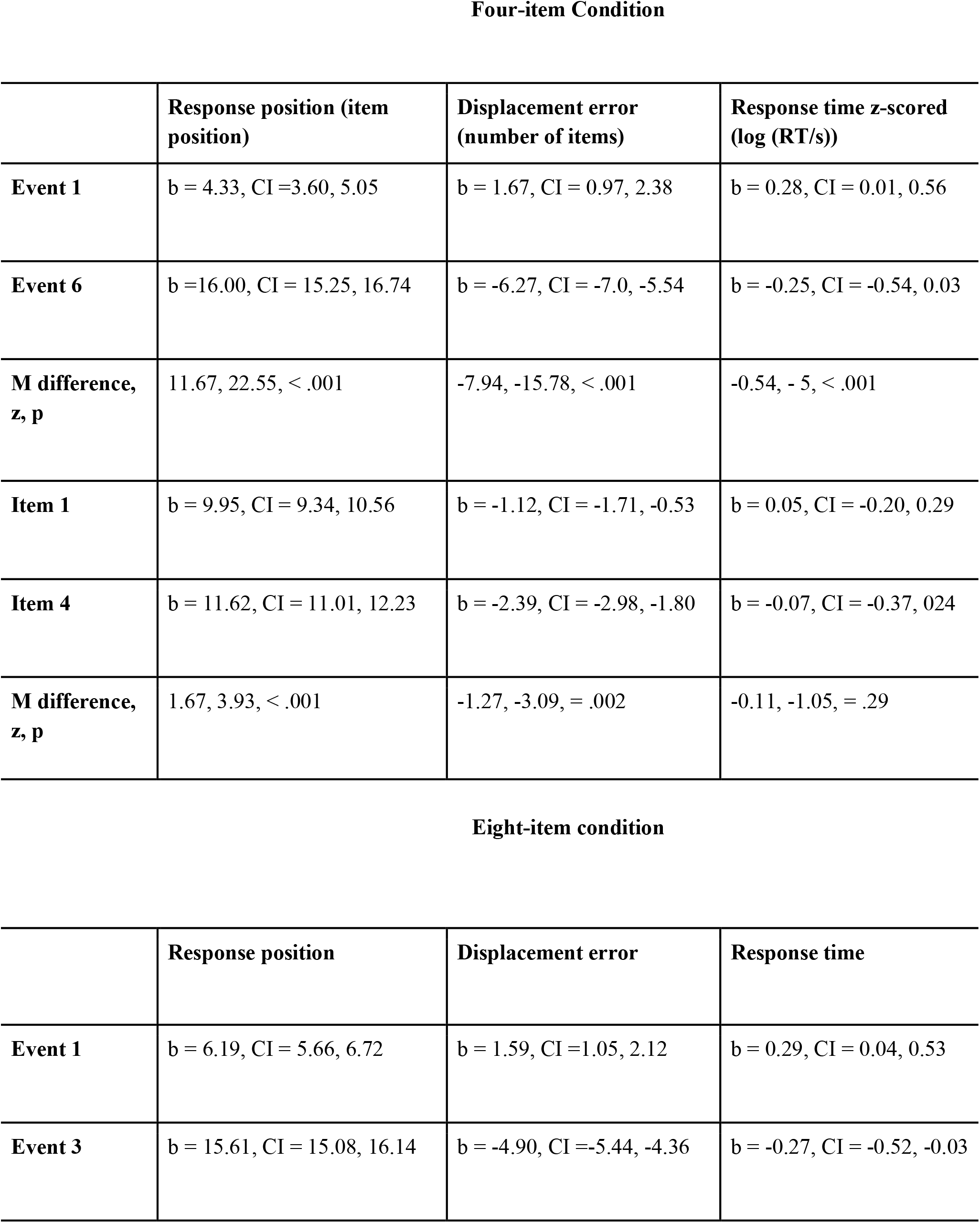
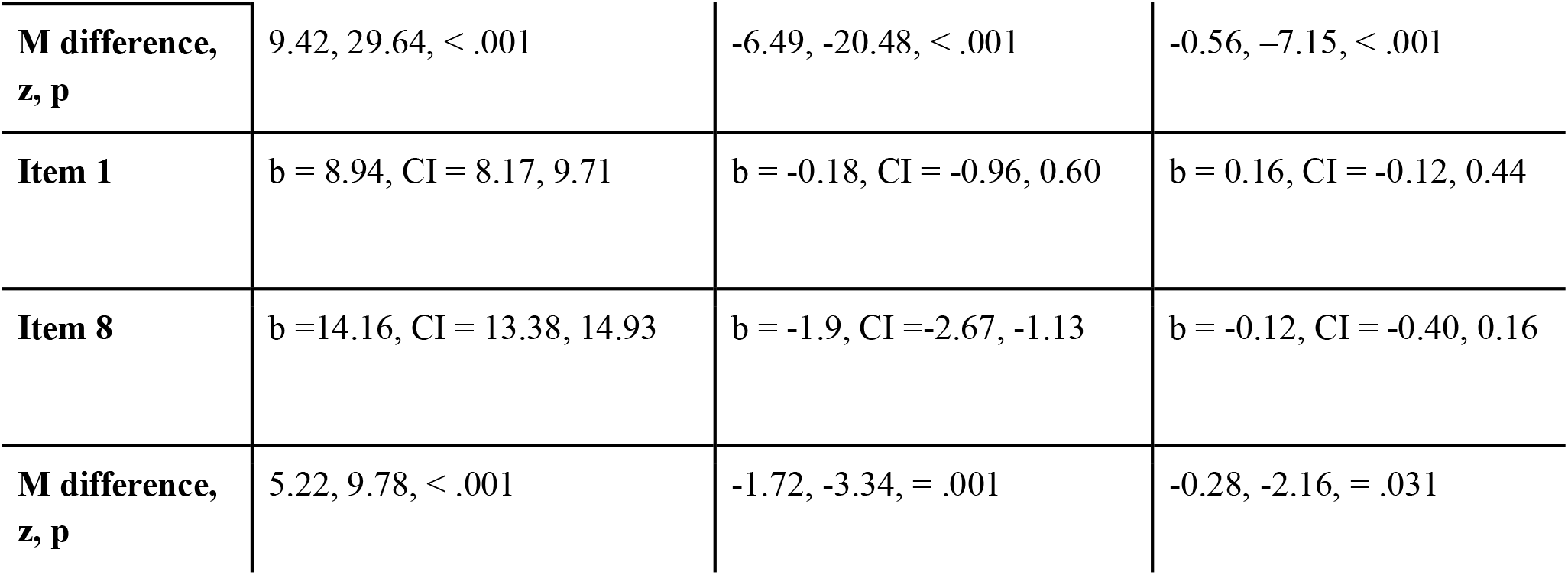
Summary of post-hoc tests of estimated marginal means for response positions, displacement errors and response times. Top: Mean differences between Event 1 and Event 6 and Item 1 and Item 4 for the four-item condition. **Bottom:** Mean differences between Event 1 and Event 3 and Item 1 and Item 8 for the eight-item condition. The eight-item condition produced significant differences in response times between the first and last items with events, whereas the four-item condition did not. M: mean difference z: z-statistic p: p-value b: beta CI: confidence interval.

Four-item Condition
|  | Response position (item position) | Displacement error (number of items) | Response time z-scored (log (RT/s)) |
| --- | --- | --- | --- |
| Event 1 | b = 4.33, CI = 3.60, 5.05 | b = 1.67, CI = 0.97, 2.38 | b = 0.28, CI = 0.01, 0.56 |
| Event 6 | b = 16.00, CI = 15.25, 16.74 | b = -6.27, CI = -7.0, -5.54 | b = -0.25, CI = -0.54, 0.03 |
| M difference, z, p | 11.67, 22.55, < .001 | -7.94, -15.78, < .001 | -0.54, -5, < .001 |
| Item 1 | b = 9.95, CI = 9.34, 10.56 | b = -1.12, CI = -1.71, -0.53 | b = 0.05, CI = -0.20, 0.29 |
| Item 4 | b = 11.62, CI = 11.01, 12.23 | b = -2.39, CI = -2.98, -1.80 | b = -0.07, CI = -0.37, 0.24 |
| M difference, z, p | 1.67, 3.93, < .001 | -1.27, -3.09, = .002 | -0.11, -1.05, = .29 |

**Table 1. Summary of post-hoc tests of estimated marginal means for response positions, displacement errors and response times.**
|  | Response position | Displacement error | Response time |
| --- | --- | --- | --- |
| Event 1 | b = 6.19, CI = 5.66, 6.72 | b = 1.59, CI = 1.05, 2.12 | b = 0.29, CI = 0.04, 0.53 |
| Event 3 | b = 15.61, CI = 15.08, 16.14 | b = -4.90, CI = -5.44, -4.36 | b = -0.27, CI = -0.52, -0.03 |
| **M difference,**<br>**z, p** | 9.42, 29.64, < .001 | -6.49, -20.48, < .001 | -0.56, -7.15, < .001 |
| **Item 1** | b = 8.94, CI = 8.17, 9.71 | b = -0.18, CI = -0.96, 0.60 | b = 0.16, CI = -0.12, 0.44 |
| **Item 8** | b = 14.16, CI = 13.38, 14.93 | b = -1.9, CI = -2.67, -1.13 | b = -0.12, CI = -0.40, 0.16 |
| **M difference,**<br>**z, p** | 5.22, 9.78, < .001 | -1.72, -3.34, = .001 | -0.28, -2.16, = .031 |

#### 3.1.4. Differences in response times based on item position

While we have characterised temporal compression as the under and over estimation of position on a timeline. The recording of response times allows us to also relate the present study to previous work on mental scanning times. We expected the number of items within events and the number of events within a study block to influence the time it takes to place an item on a timeline. Our prior sections described the effects for the four-item and eight-item separately. We now compare the differences in response times across both conditions. The response time metric is taken from previous work on time to replay navigational routes ^24^ which calculates a ratio of the number of seconds to mentally replay a navigation route to the number of seconds of the original presentation. While previous studies have focused on the time to mental replay spatial navigation, here we specifically examine the difference in response times based on the position of an item in the sequence, that is whether an image occurred in the first or last item position.

For the four-item condition response time ratios were calculated by dividing the number of seconds to present the sequences (64.8) by the differences in mean log-transformed, z-scored response times between the first item in the first event and the last item in the last event (M = −0.63, SD = 1.85). The four-item condition produced a factor of 103:1. For the eight-item condition dividing the number of seconds to present the sequences (64.8) by the differences in mean response times between the first item in the first event and the last item in the last event (M = 1.44, SD = 0.80). The eight-item condition produced a ratio of 44.97:1.

These results show that presenting four-items per event in six events produced a smaller difference in response times between the beginning and end of the sequence than the eight-item condition. While both sequences took the same total number of seconds, fewer items within events and more boundaries may allow for more ‘skipping’ to the closest event boundary ^17^. This first observation suggests that the highest level of event structures (here, rooms) impact the time to identify the timeline position in memory at the lowest level (here, item). Notably, previous studies have identified response time ratios in rodents between 6 and 64 ^42^. The present results suggest that humans are capable of higher compression factors (44 to 103) and variations in response time based on item and boundary positions may inform temporal displacement errors in memory for ‘when’.

#### 3.1.5. The impact of event boundary and number of items on temporal distances

Following previous work ^31^ we analysed the mean distances between temporal positions based on the position of items within or across events. Herein, if event boundaries (room transitions) provided a ‘reset’ of context, item pairs occurring on either side of a boundary should be placed closer together than pairs occurring within an event (room). To the contrary, if boundaries provide an acceleration of context drift ^30^, items occurring on either side of a boundary should be placed further apart than items occurring within an event.

For the four-item condition (Figure 4, bottom), a Bayesian paired sample t-test between pair types (Across early, Across late) provided moderate evidence against an effect of temporal distance between Across early (M = 1.75, CI = 0.03, 3.47) and Across late (M = 1.32, CI = −0.56, 3.20), (BF10 = 0.20). For the Bayesian analysis, CI is the credibility interval. A Bayesian paired sample t-test between pair types (Across late, Across) provided anecdotal evidence for an effect of temporal distance between Across late (M = 1.32, CI = −0.56, 3.20) and Across (M = 1.90, CI = 0.44, 3.37), (BF10 = −.43).

**Figure 4:**
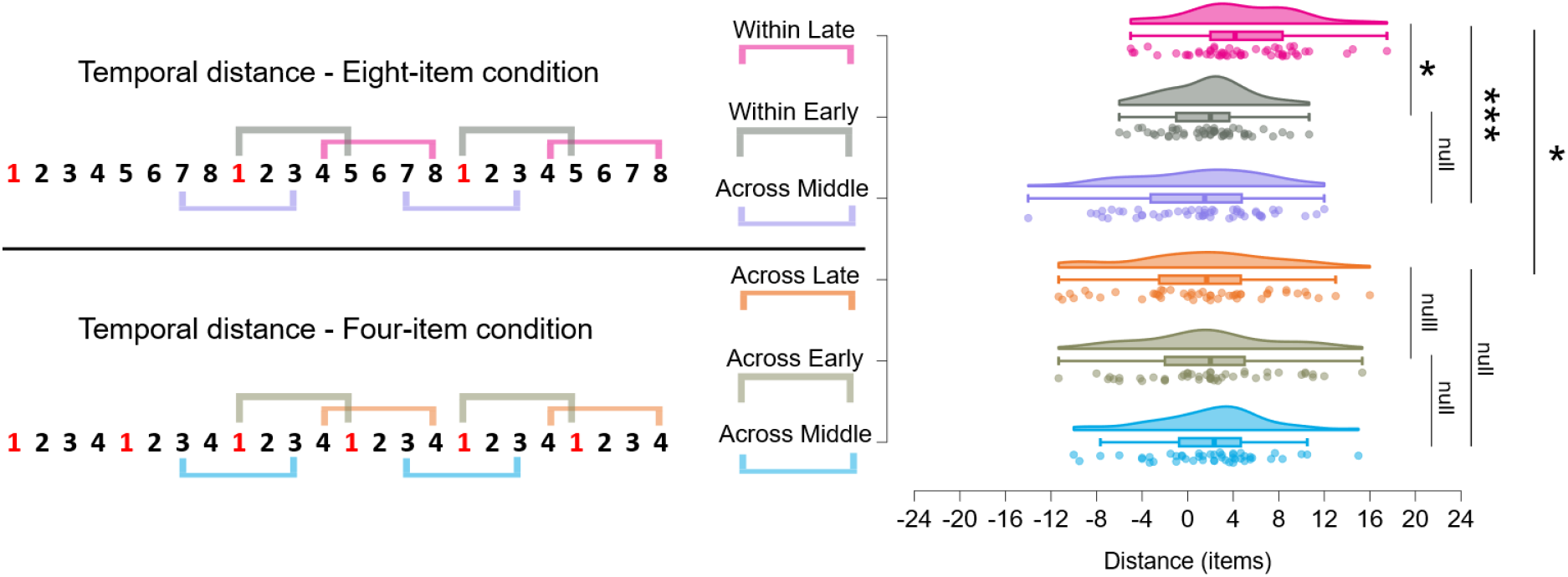
Temporal distances between item pairs within or across events. **Bottom**: Raincloud plots (including violin and boxplots) of temporal distances between pairs of items from the four-item condition. The analysis was carried out on item pairs separated by three items and an event boundary (Across) or separated by three items and an event boundary early in an event or late in an event (Across early, Across late). **Top**: Raincloud plots of distances between pairs of items from the eight-item condition for item pairs separated by three items from the start or from the end of an event (Within early and Within late) or separated by three items and an event boundary (Across). Note that the distances for the Within late pair in the Eight-item condition were larger than for the Within early and the Across pair. The distances between the Within late pair in the Eight item condition were also larger than the distances between the Across late pair in the Four-item condition. Pairs crossing a boundary in both conditions were placed closer together than the within late pair in the eight-item condition. Negative distances indicate an incorrect temporal order, with the last item in a pair placed earlier than the first item. Error bars represent Bayesian Credibility Intervals (CI).

For the eight-item condition (Figure 4), Bayesian paired samples t-test between pair types (Within Early versus Within Late) provided moderate evidence in favour of an effect of temporal distance between Within Early (M = 1.71, CI = 0.70, 2.72) and Within Late (M = 4.68, CI = 3.32, 6.04), (BF10 = 9.87). Bayesian paired samples t-test between pair types (Within Late versus Across) provided extremely strong evidence in favour of an effect of temporal distance between Within Late (M = 4.68, CI = 3.32, 6.04) and Across (M = 0.95, CI = −0.63, 2.52), (BF10 = 100.25). Bayesian paired samples t-test between pair types (Within Early versus Across) provided moderate evidence against an effect of temporal distance between Within Early (M = 1.71, CI = 0.70, 2.72) and Across (M = 0.95, CI = −0.63, 2.52), (BF10 = 0.20). For the eight-item condition, pairs occurring late within an event were placed further apart than pairs occurring early within an event or across events.

Comparison of distance between the ‘across late’ pair in the four-item condition and the ‘within late’ pair of the eight-item condition revealed strong evidence in favour of a larger temporal distance for the eight-item condition (M = 4.68, CI =3.32, 6.04) than the four-item condition (M = 1.32, CI = −0.56, 3.20) (BF10 = 11.41). The ‘across late’ pair occurring on either side of a boundary in the four-item condition (Figure 4) resulted in smaller temporal distance than the equivalent pair occurring within boundaries in the eight-item condition (Figure 4).

The larger temporal distance between the late pairs within an event suggests that an acceleration of contextual drift occurs as information accumulates. Such an acceleration of contextual drift can result in a greater dissimilarity of context, which in turn, may determine the larger temporal distances between late pairs than early pairs. The result also supports the account of each event boundary providing a ‘reset’, so that item pairs across boundaries are placed closer together than the late pairs within boundaries.

Both the four-item and eight-item condition produced an overall temporal compression effect (see Figure 3). However, when increasing the quantity of information within events, such as in the eight-item condition, the compression did not apply equally to every item. The ‘within late’ pairs were placed further apart than the early pairs within events (Figure 4). The larger temporal distance for the late pairs within events was due to fewer negative distances than the early pairs within events. As negative distances indicate an incorrect temporal order, memory for temporal order was impaired following an event boundary but improved for pairs of items occurring at the end of eight-item events. The results are consistent with a ‘resetting’ ^31^. However, the results could also be due to an acceleration of context drift increasing temporal distances between items as information accumulates within an event.

### 3.2. Memory for ‘where’. Do nested event structures influence the accuracy of spatial memory?

Analysis of temporal response positions, displacement errors and response times revealed extremely strong evidence for the influence of nested event structures on the temporal compression and temporal distances in memory for when. We also questioned whether the event structure would influence memory for ‘where’. Would participants be able to correctly place an item in the spatial location on the grid? Do nested event structures influence memory for ‘when’ independently from memory for ‘where’?

#### 3.2.1. Do participants perform better than chance in memory for ‘where’?

For spatial responses, performance was first evaluated in terms of proportion correct relative to chance. For a 3-by-3 grid, the chance level was 0.11 (1/9) for each item. The analysis revealed extremely strong evidence in favour of a better than chance performance for the four-item condition (M = 0.16, CI = 0.14, 0.18, BF = 62380) and extremely strong evidence in favour of a better than chance performance for the eight-item condition (M = 0.18, CI = 0.16, 0.20, BF = 1.50x10^11^). Bayesian paired sample t-tests revealed only anecdotal evidence in favour of a higher proportion correct for the eight-item condition (BF10 = 0.33). Participants showed a better than chance performance for placing items in the correct spatial position. We then questioned whether performance would vary based on the position of an item within an event sequence.

#### 3.2.2. Nested event structure affects the ‘where’ of memory

Correct spatial responses were submitted to linear mixed models following the same approach as the analysis of memory for ‘when’ with dependent variables of proportion correct for both the four-item and eight-item condition. The model included fixed effects of event (1-6 for the four-item condition and 1-3 for the eight-item condition) and item position (1-4 for the four-item position and 1-8 for the eight-item condition) a random effect grouping factor of participant (1-26) with random intercepts. Models were evaluated based on Akaike information criteria (AIC).

For the four-item condition, the analysis included 602 observations. The model including fixed effects of event position alone (log likelihood = 71.18, AIC = −126) outperformed a model containing fixed effects of both event and item position (log likelihood = 72.51, AIC = −123) and a model containing an interaction between event and item position (log likelihood = 76.18, AIC = −100). The analysis revealed no evidence for an effect of event position (χ^2^(5) = 7.94, p = .16) on the proportion of correct ‘where’ judgements. Fixed effect estimates identified a grand mean for proportion of correct responses as (b = 0.16, SE = 0.012, t (26.13) = 13.35, p < .001). The post-hoc t-tests between first and last events showed a decrease from Event 1 (b = 0.18, CI = 0.14, 0.23) to Event 3 (b = 0.11, CI = 0.07, 0.16) with an estimated mean difference of −0.07 (z = −2.34, p = .019, CI = −0.127, −0.01). The results suggest that participants performed better for items located in the first events than items located in the middle of the sequence. The temporal compression to the middle of the sequence may increase the confusability of spatial positions for items within the middle event while the spatial positions for the items in the first events benefitted from being closer to the boundary at the beginning of the sequence.

For the eight-item condition, the analysis included 624 observations. The model containing fixed effects of both event and item position (log likelihood = 57.1, AIC = −90.20) slightly outperformed a model including only event position (log likelihood = 49.9, AIC = −89.81) and a model containing an interaction between event and item position (log likelihood = 63.13, AIC = −74.25). The analysis revealed evidence in favour of an effect of item position, (χ^2^(7) = 14.39, p = .045) but not event position (χ^2^(2) = 0.83, p = .66) on the proportion of correct ‘where’ responses. Fixed effect estimates identified a grand mean for proportion of correct responses (b = 0.18, SE = 0.01, t (26.00) = 13.02, p < .001). The eight-item condition showed an effect of the position of an item within an event but no difference in overall performance in the proportion of correct responses for each event.

The results support the initial prediction of memory for ‘when’ varying independently from the memory for ‘where’. Interestingly, in the four-item condition, Event 1 showed an improved performance in comparison to the chance performance of Event 3. The temporal compression around the midpoint of the sequence may have made it harder to reconstruct the locations of items on the grid within an event.

#### 3.2.3 Does memory for ‘where’ require the scanning of a timeline?

To determine if the memory for ‘where’ also required a scanning of a timeline to access the information within an event we questioned whether the positions of an item within an event sequence would influence the speed of responses. Do participants first scan through the timeline in memory before placing an item in a spatial position within a room? We followed the same approach as the previous analysis of response times by removing response times that were greater than 2.5 times the standard deviation of the mean then log transforming and z-scoring the remaining response times and submitting the response times to linear mixed models.

For the four-item condition the analysis included 594 observations. The model including fixed effects of event position alone (log likelihood = −730, AIC = 1476) outperformed a model containing fixed effects of both event and item position (log likelihood = −729, AIC = 1482) and a model containing an interaction between event and item position (log likelihood = −717, AIC = 1488). The analysis revealed evidence in favour of an effect of event position, (χ^2^(5) = 11.43, p = .044) on response times for placing an item in the correct spatial position on the grid. Fixed effect estimates identified a grand mean for proportion of correct responses as (b = 3.652×10^-4^, SE = 0.13, t (26.01) = −0.003, p = 0.998).

For the eight-item condition the analysis included 589 observations. The model including fixed effects of event position alone (log likelihood = −748, AIC = 1507) outperformed a model containing a fixed effect of both event and item positions (log likelihood = −745, AIC = 1515) and a model containing an interaction between event and item position (log likelihood = −741, AIC = 1534). The analysis revealed no evidence for an effect of event position, (χ^2^(2) = 0.46, p = .80) on memory for ‘where’. Fixed effect estimates identified a grand mean for transformed response times (b = 0.001, SE = 0.12, t (25.93) = 0.06, p = .95).

The analysis did provide evidence in favour of a difference in response times for remembering where an image appeared for the four-item condition. However, no differences in response times were found for the eight-item condition. Additionally, the eight-item condition produced a better than chance performance for identifying where an image appeared. The results are consistent with requiring a decompression of encoded information to access details about the contents of an event. A greater compression for the four-item condition (Figure 4) resulted in a difference in response times for the correct identification of ‘where’ which also increased the difficulty in distinguishing the ‘where’. The results are consistent with the difference in proportion of correct responses in the four-item condition (section 3.2.2).

### 3.3. Memory for ‘what’. Does the number of items within events influence recognition?

Memory performance can improve when reducing the quantity of information within events^33,43^. Thus, we questioned whether recognition performance would improve when presenting fewer items within events (four *vs.* eight items).

#### 3.3.1. Recognition performance based on the number of items within an event

We analysed differences in recognition performance based on presenting four items per event or eight items per event. Recognition performance was analysed with d’ prime between the four and eight-item conditions (Figure 5A). Bayesian t-tests were conducted to compare d prime scores for the four and eight-item conditions.

**Figure 5:**
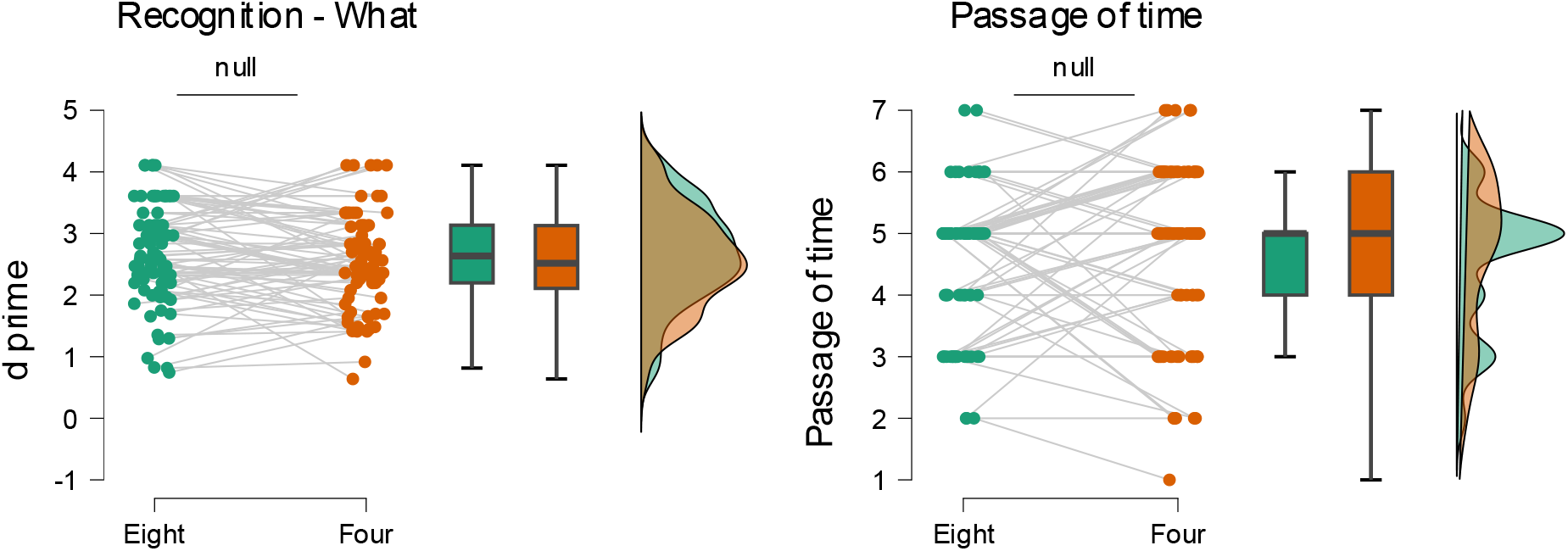
Recognition performance and passage of time judgements. **Panel A**: Raincloud plots comparing recognition performance (d prime) between the eight-item condition (green) and the four-item condition (orange). Box plots and half-violin plots highlight no difference in recognition performance or passage of time judgements between the four and eight-item condition.

The analysis revealed anecdotal evidence in favour of a null effect between the four-item (M = 2.55, CI = 2.38, 2.73) and the eight-item condition (M= 2.69, CI = 2.5, 2.89) (BF = 0.39). Participants performed extremely well in correctly distinguishing old and new images with no influence of event structure on recognition performance.

The event structures did not influence either recognition performance. Given recognition performance was high in both conditions (Figure 5) eight images per room may not have sufficiently overloaded working memory to drive a drop in recognition performance. The result highlights the differences in ‘memory for ‘when’ (Figure 2) without an associated difference in memory for ‘what’.

#### 3.3.1. Proportion correct in memory for ‘what’

We also aimed to identify whether the ability to correctly identify an image would vary based on the position of the item in the event and the position of the event in the sequence. The proportion of correctly recognised images were submitted to linear mixed models following the same approach as the analyses for the proportion of correct responses in memory for ‘where’. Would participants show improved performance for correctly identifying images in the first and last positions?

The proportion of correct recognition responses (hits) were submitted to linear mixed models following the same approach as the analysis of memory for ‘where’ with dependent variables of proportion correct for both the four-item and eight-item condition.

For the four-item condition the analysis included 624 observations. The model including fixed effects of event position alone (log likelihood = −22.58, AIC = 61.15 outperformed a model containing fixed effects of both event and item position (log likelihood = −22.48, AIC = 66.97) and a model containing an interaction between event and item position (log likelihood = −12.45, AIC = 76.9). The analysis revealed no evidence for an effect of event position, (χ^2^(5) = 6.07, p = .30) on the correct recognition of an image. Fixed effect estimates identified a grand mean for proportion of hits as (b =0.74, SE = 0.04, t (26.00) = 21.20, p < .001).

For the eight-item condition the analysis included 624 observations from 26 participants. The model containing fixed effects of only event position (log likelihood = 26.45, AIC = −42.90) outperformed a model containing fixed effects of both event and item position (log likelihood = 29.52, AIC = −35.03) and a model containing an interaction between event and item position (log likelihood = 40.94, AIC = −29.87). The analysis revealed no evidence for an effect of event position, (χ^2^(2) = 4.30, p = .116) on the correct recognition of an image. Fixed effect estimates identified a grand mean for proportion of correct responses (b = 0.76, SE = 0.03, t (26.00) = 29.71, p < .001).

Memory for ‘what’ did not vary based on event or item positions. In both eight-item and four-item events, participants show a consistently high performance (>70%) for identifying images.

#### 3.3.2. Response times in memory for ‘what’

We also questioned whether correctly identifying an image would require a memory scan of timeline with slower response times for first events as found in the analysis in memory for ‘when’. Consequently, we took response times for correctly recognised images, removed those that exceeded 2.5 standard deviations greater than the mean, log transformed and z-scored the response times for ‘hits’ and conducted linear mixed models.

For the four-item condition the analysis included 595 observations. The model containing fixed effects of both event and item position (log likelihood = −800, AIC = 1623) slightly outperformed a model containing fixed effects of event position alone (log likelihood = −804, AIC = 1625) and a model containing an interaction between event and item position (log likelihood = −789, AIC = 1632). The analysis revealed evidence in favour of a difference in response times for item position, (χ^2^(3) = 8.55, p = .036) but no evidence of event position, (χ^2^(5) = 4.10, p = .54) for the correct recognition of an image. Fixed effect estimates identified a grand mean for proportion of correct responses as (b = −0.01, SE = 0.09, t (26.03) = −0.068, p = 0.95) and a significant effect for item 3 (b = −0.18, SE = 0.06, t (569.33) = −2.77, p = .01). The analysis did not provide evidence in favour of a difference in response times between the first and last events, memory for ‘what’ did not require a scanning of memory from the last event to the first event. However, the significant difference in response times for items within events suggests that identifying the ‘what’ requires more time to decompress items within events than identifying the correct effect.

For the eight-item condition the analysis included 621 observations from 26 participants. The model containing fixed effects of event position alone (log likelihood = −821, AIC = 1652) outperformed a model containing a model with fixed effect of both event and item positions (log likelihood = −816, AIC = 1657) and a model containing an interaction between event and item position (log likelihood = −814, AIC = 1680) . The analysis revealed no evidence for an effect of event position, (χ^2^(2) = 0.51, p = .78) on the correct recognition of an image. Fixed effect estimates identified a grand mean for transformed response times (b = −0.01, SE = 0.10, t (26.02) = −0.08, p = .94).

Altogether, the analysis of response times suggests that scanning a timeline of events in memory is not required when explicitly asked to recognise an image.

### 3.4. Passage of time

Previous studies have identified an expansion of recollected time when experiencing contextual-changes provided by turning corners ^25^ or when increasing the number of events experienced ^44,45^. In the present study, both conditions presented the same number of images within the same number of seconds. Hence, we expected to find differences in passage of time judgements based on the event structure (eight items in three events versus four items in six events). Specifically, we asked whether increasing the number of events (from three to six) would result in faster passage of time judgements.

We analysed differences in passage of time Judgements based on the number of events experienced (six or three) (Figure 5B). Bayesian paired samples t-tests revealed moderate evidence against an effect of condition between the four-item (M = 4.71, CI = 4.40, 5.04) and eight-item condition (M = 4.58, CI = 4.32, 4.83), (BF = 0.19). The different number of events (6 or 3) did not impact passage of time judgements. The event structures did not influence passage of time judgements. The results suggest that the passage of time judgments may have depended on the memory for ‘what’, in terms of how many items could be remembered rather than requiring a scan of a timeline of events. However this interpretation may be specific to the experimental design aimed at testing memory. Future work on passage of time judgments may benefit from asking participants to provide a record of their internally generated thoughts in addition to passage of time estimates and duration estimates ^45,46^ .

## 5. General Discussion

In this study, we aimed to determine how nested event structures influence the relationships between the what, when, and where of memory. Our main finding is that the quantity of information embedded within nested event structures yields a hierarchy of temporal compression for the when of memory, independently of the what and where of memory. We will first address how our results demonstrate a nested compression (Fig. 6) in memory for ‘when’. Specifically, how the temporal distances between pairs of items occurring within or across events depend on the number of items within events (either 4 or 8 per room). Notably, the strong temporal compression effects in memory for ‘when’ were not found in memory for ‘what’ or ‘where’. We will then highlight how differences in response times identifying when an image appears require backward scanning of the sequence, in contrast to previous work that has suggested a forward scanning of memory. Last, we discuss each observation and discuss their implication for existing models and suggest an alternative based on “nested compression” (Fig. 6).

**Figure 6:**
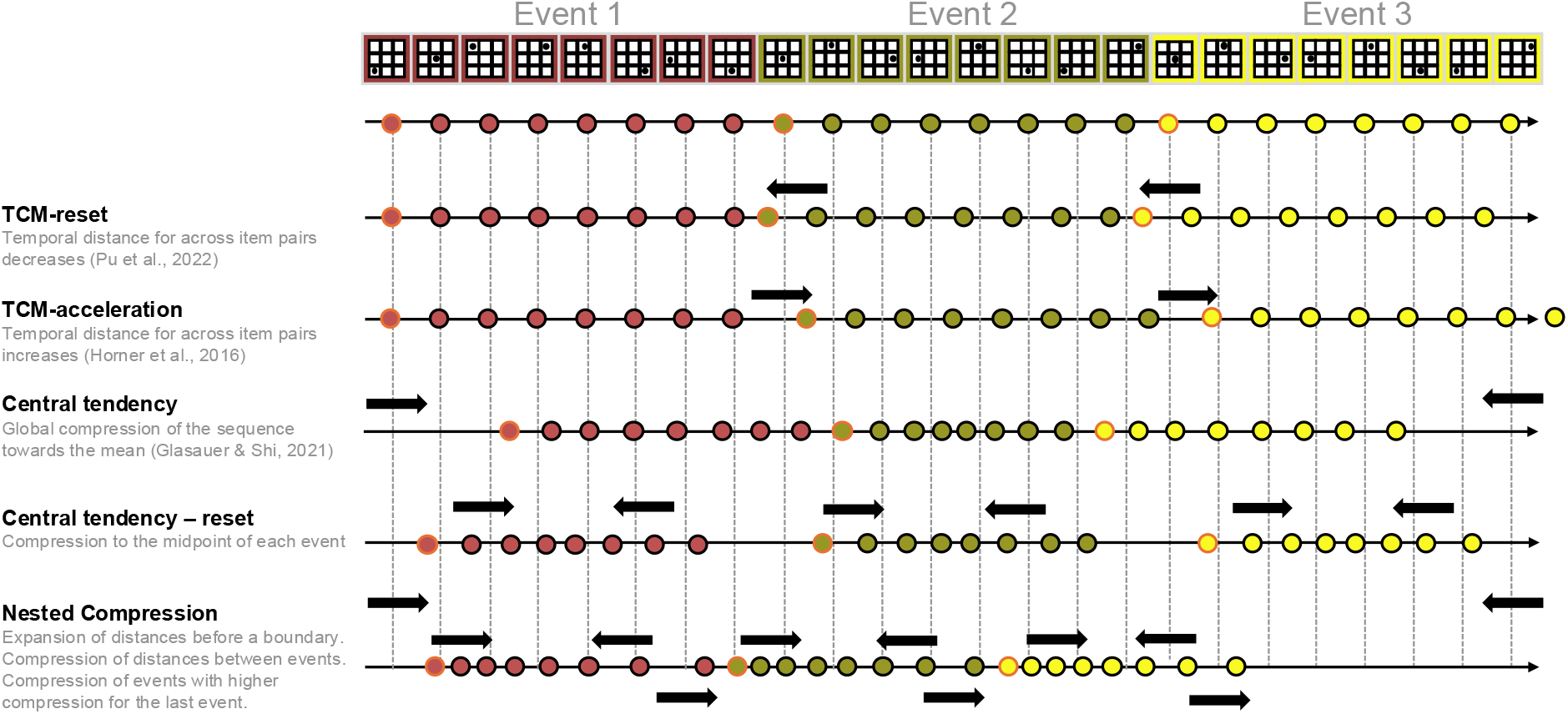
Theoretical predictions of temporal distortions in memory as captured by temporal distances. As explained in text, TCM models (1st and 2nd rows) cannot fully explain the data. Here, we updated Figure 1 with three additional models to try and account for our observations. In the 3rd and 4th rows, we attempt to predict central tendency effects typically reported in timing research. In the 3rd row, the central tendency applies to the full sequence of events, which would predict a regression to the mean over the entire duration of the trial (always 64.8s) yielding a global compression effect. This is not what we observed. In this experiment, all durations were maintained at the item, event, and thus at the full sequence level, making no contextual differences for predictions of Central Tendency changes. Hence, if we entertain the possibility of reset in the sequence so that durations are discretized and computed per event (4th row), the central tendency would now be applied by event with a compression effect within the event, but an expansion effect across events (see arrows). Again, this is not what we observed. Hence, we conclude that existing central tendency effects are insufficient to account for the distortions in temporal memory observed here. The alternative in the last row, Nested Compression, is detailed in text and will be detailed in subsequent dedicated paper.

### A hierarchy of nested temporal compression

We questioned how the temporal distances between memories depend on the structure of events and found strong compression effects that occurred at both the level of items within events and events within sequences. Crucially, the rate of compression depended on the structure of information.

The analysis of temporal distances for the four-item condition revealed no differences between the within and the across item pairs. This observation is unexpected and inconsistent with prior suggestions that boundaries increase temporal distances by providing an acceleration of context drift ^30^ (Fig. 1; Fig. 6, TCM-acceleration) In the present study, room transitions provided predictable contextual shifts. The absence of doorways may have allowed for more consistent reconstruction of temporal positions as compared to another study, in which contextual shifts were prediction errors between rooms separated by doorways ^30^. We would expect to see similar nested compression effects if the present study were re-run with doorways between locations. However, the temporal distances between pairs occurring on either side of a boundary may depend on a graded magnitude of prediction error, which would be larger with doorways than without.

While the four-item condition showed no differences between pairs occurring within or across boundaries, we did find differences in the eight-item condition. When presenting eight-item per event, pairs occurring late within an event were placed further apart than either across-item pairs or within-item pairs occurring early within an event. The greater temporal distances for later pairs within eight-item events in comparison to early pairs and across event pairs suggest that the rate of drift may accelerate as information accumulates. The smaller distances between items occurring at the start of an event is consistent with the reinstatement of a start of event context ^31^. However, the resetting of temporal distances at the start of an event could instead be described as a clearing of information acting to reset the rate of drift, consistent with event segmentation theory ^6^. Identifying the temporal distances between later pairs within events requires additional reconstruction of contextual information with a faster drift rate, increasing temporal distances. The quantity of information within events influences the temporal distances between items in memory.

The larger distances between late items are also consistent with recent work finding an expansion for duration estimations for the endings of events ^47^. In the present study, the ‘within late’ comparison could be described as depending on an acceleration of context drift as information accumulates, resulting in greater contextual dissimilarity and greater temporal distances (Fig. 6, TCM-acceleration). Notably, in the present study, participants were presented with a single item and had a free choice of placement on a timeline, whereas previous studies presented specific pairs and asked for judgements of temporal order (which came first) ^30,31^. Increasing the quantity of information within events increased the temporal distances between items in memory. From the perspective of memory as a creative construction, the expansion of later pairs is due to the compression, decompression, and imperfect reconstruction of context. Instead of acceleration and a reinstatement of temporal context at boundaries, the resetting of temporal distances may be better described by a positional coding account ^48^. The compression of information then also provides structure from which temporal order and predictions may be computed from a difference in distances.

For the comparison of distances between pairs of images, shorter distances also included more negative scores, indicating an incorrect temporal order. The larger distances for later pairs within eight-item events indicate an improved memory for the temporal order. The results are consistent with previous findings showing greater confusability for across event pairs ^49^. However, in the present study there was an improvement in temporal order memory for late pairs due to larger temporal distance between items. One possible explanation for the differences in confusability within and across events is an acceleration of the rate of context drift driving greater dissimilarity between items which in turn improves memory for temporal order. When presenting four images per room, there were no differences in temporal distances between pairs occurring within or across an event. However, when presenting eight images per room, late pairs were placed further apart than either early pairs or pairs occurring across an event boundary. The contrasting results on how boundaries affect temporal order memory points to a flexibility in the processes involved in judgements of ‘when’. Performance will vary based on the quantity of information within an event, nested event structures, the type of boundaries employed such as changes in contextual stability ^12,50^ versus prediction errors ^14^. Performance will also vary with the availability of narrative schema and how memories are tested ^51,52^. For example, removing the end of a movie breaks the narrative structure and results in increasing temporal displacement errors of reconstructed events that occur towards the end of the cut movie ^53^. Importantly, schema can incorporate new information, but the updating of schema may only occur if sufficient contextual change has occurred. Information within events is compressed and encoded into long-term memory when sufficient change occurs.

### Nested compression rate and scaling: backward versus forward scanning

The four-item condition showed a greater overall compression rate at the level of events within study blocks, suggesting different compression rates depending on the temporal scale of interest. However, both conditions showed a significant difference in response times between the first and the last item in the sequence. Additionally, slower response times in the ‘when’ of memory for both the first items within events (for the eight-item condition) and the beginning and end of the sequence suggest that participants scan backwards from the end ^54^. Our results contrast with the forward scanning demonstrated by the stepping stone model ^17^, but are consistent with a memory scan that seeks to employ boundaries as temporal landmarks ^55–57^ and skip to the closest event boundary. Rather than an exhaustive ‘scanning’ of memory, participants skip backwards to an event boundary and then reconstruct information starting from the last item within an event. However, the memory system may flexibly adapt to a task based on the quantity of information and the rate of compression at the temporal scale of interest. While the present study employed a tightly controlled virtual environment, events arise more naturally in everyday life from changes that are meaningful to an individual. In the present study, when presenting random sequences of images, participants scan backwards from the present. However, when meaningful narratives are employed such as in movies or the telling of stories participants may scan forward from the beginning of the story ^52^.

### Independence of when from where and what

In this experiment, we used the movement transitions between virtual rooms as boundaries ^33,43^. While spatial gaps between events in adjacent rooms were varied, the number of seconds was maintained identical in all conditions. The current study is consistent with the dependency of temporal experience on the rate of drift within events at nested timescales ^7,55,58–60^. In this study, we found that the temporal displacement errors consistently increased with the distance of items from the start of the event sequence, with overestimations for early items, and consistently increasing underestimations for items later within an event and later within a study block. The results support the view that the compression and reconstructive processes of memory within nested events drives the experience of temporal displacement.

Importantly, while the present study employed spatial transitions to provide boundaries, spatial transitions are not the only means of defining the start and end of events, nor unique in providing temporal displacement effects. A recent study demonstrated reduced temporal displacements for items occurring at musical or emotional transitions, which define boundaries in terms of perceptual, valence and arousal based, context shifts ^41^. We would predict the same pattern of results reported in the present study to be found when employing non-spatial boundaries. Additionally, recent work suggests that transient Bayesian surprise provides a better predictor of segmentation behaviour than singular prediction errors ^13^. While a single highly salient context shift, such as moving to a new location, can influence compression and encoding, encoding may occur when an accumulation of change reaches a threshold. Spatial structure provides an effective means of controlling the presentation of events; however, temporal distances and the arrow of time rely on contextual changes and cognitive processes that are distinct from spatial processing ^61^.

Despite differences in temporal displacement errors and response times, no differences of recognition performance (what) were found, which is consistent with reconstructive processes of memory ^5,62^. The compression and reconstruction of memories allow flexible adaptation to maintain memory performance for ‘what and where’ despite variability in temporal compression (‘when’) based on the quantity of information within an event. Previous studies have demonstrated a ‘lossy’ compression process: if too much information exists between boundaries, information is lost in the compression process ^18^.

In the present study, the working memory load between boundaries (four-item versus eight-item) did not result in differences in the ability to distinguish old and new items measured by d’ or differences in the reported speed of the passage of time. Recent work has shown that an increase in the ability to distinguish old and new items produces an underestimation of time passing ^63^. The absence of differences in recognition performance in the present study are still consistent with the previous result showing a link between recognition performance and passage of time. While the conditions in the present experiment were not sufficiently distinct to drive differences in recognition performance, the strong effects of event structures on temporal compression provide a rich foundation for future work. Adjusting the quantity of information within nested events has been shown to influence recall performance ^33^, suggesting that it may be possible to adjust the quantity of information within events such to drive differences in both recognition performance and the passage of time. Remembering is a creative reconstruction that requires the decompression and reconstruction of events, which can vary in accuracy and precision of memory for ‘when’ even with equivalent recognition performance. Our results suggest that finer-grained testing of duration or passage of time judgements of different nested event structures may reveal scale-invariant effects at item or event levels ^64,65^.

### Limitations of current models and proposal

The current dominant accounts of memory for temporal order and distance are based on models of temporal context drift ^29–31^ (Fig. 1; Fig. 6 top rows) These models predict that distance effects on the basis of drift and boundaries drive changes in the similarity of context vectors, not on the basis of the quantity of information. To the contrary, our findings suggest that current models need to account for the quantity of information within events influencing compression rates at multiple nested scales, and not only on item-to-item drift rates or boundary effects, as current models predict ^29–31^. For instance, we found an expansion of temporal distances occurring before the boundary for eight-item events and a compression of temporal distances between events. The expansion of temporal distances at the end of events depended on the quantity of information (the four-item events did not produce a difference in temporal distances). The compression, in terms of both temporal distances based on item positions and response times, varied depending on the event position. We found the temporal compression for items within the last events to be larger than for items within the first events. Hence, the temporal distances between items depend on when pairs of items occur in a sequence (beginning, middle, or end) and not just if the pairs appear within or across an event boundary. We also found interactions between event and item positions for response positions, displacement errors, and response times when presenting eight items within three events, but not when presenting four items within six events. Hence, neither the resetting nor the accelerating variants of temporal context models (Figure 6 top rows) can fully explain these observations.

We found an expansion of distances for pairs occurring before a boundary, and a reset of distances for pairs occurring after a boundary along with a compression of events which increased for events later in the sequence. We also found a compression of events, which increased for later events with the largest underestimation of temporal position for items within the last event. The result suggests that a nested compression of information underlies memory for ‘when’ (Fig. 6, last row). While we interpret the differences between the four and eight-item conditions in terms of the quantity of information within events, an alternative interpretation is a buildup of proactive interference ^66^. However, if experiencing event boundaries clears information from working memory, a buildup of proactive interference could be described as an increase in entropy as information accumulates within working memory ^13^. The higher the entropy (possible configurations of information), the higher the proactive interference. Experiencing an event boundary clears the information from working memory, which ‘resets’ the buildup of proactive interference. Future work could investigate the effect of nested event structure on the entropy in memory for ‘when’.

While the present study was focused on examining memory for ‘when’ based on how event structures influenced temporal distances based on changes in temporal context, the results could also be interpreted in terms of central tendency effects of timing (Figure 6, central tendency). That is, rather than a positional account of temporal distances based on the dissimilarity of temporal context, placing an item on a timeline could be achieved by reproducing a duration. For example, instead of remembering the first item in the second event, participants could make a duration estimate to say that the presented image occurred after approximately ten seconds.

The central tendency effect is typically not studied in the context of temporal structures or nestedness. It has been studied in the context of Bayesian timing models with some contextual effects. One important study showed that a given objective duration will be overestimated/dilated or underestimated/compressed according to the temporal context in which it is being tested ^67^. In these studies, context is understood as the set of stimulus durations that is being presented or produced (i.e. larger/medium/shorter duration prior contexts, respectively); in our study, temporal context refers to the temporal distances between items and events i.e. the similarity of context vectors. Crucially, in our study, we do not change the durations between items, equivalent durations were embedded within a nested structure. Hence, objectively speaking, there is no change of temporal context seen from the perspective of a Bayesian model uniquely considering priors in terms of magnitudes or durations (let’s call this scalar timing); however, by using a sequence of rooms, we essentially build a situational context in which we predict resetting between events (Fig. 6; central tendency - reset, speculative working hypothesis).

The resetting between events is itself predicted to induce a temporal context drift, which impacts the temporal distance between items and events. Note however, that even if we presented nothing, a temporal context still drifts ^29^. Hence, the difference in event structure changes the rate of the drift and based on the TCM models a different context vector is associated with each image in the sequence. Let’s call this the temporal context view. Hence, in our experimental design, the scalar timing is neither changed at the item level, nor at the event level (neither the duration, nor the number of items is varied across conditions). The temporal context is expected to change at both levels. If memory for when depends on reproducing the durations between items, then we would expect a pure central tendency effect with both early and late items pulled towards the mean of the sequence. Hence, our prediction of central tendency without considering the temporal context would simply predict a regression to the mean of the duration we used (Figure 6, central tendency).

However, we can now entertain elaborating on the prediction of the central tendency in our experimental design incorporating the notion of temporal context (Figure 6, central tendency-reset). This requires several assumptions that have not previously been made in the literature. First, we need to posit that the scalar property of temporal memory (that is, the actual duration between items) is transposed onto the timeline that participants used to provide their “when” positioning response. In other words, we need to take as granted that time is spatialized in a way that the positioning on the timeline represents a scalar magnitude value ^68^ and not simply an event position. Second, we need to consider that participants’ response strategy on this timeline would have an initial starting point, which could be the very beginning of the timeline (i.e. the very first item in the sequence). While this is intuitive and taken as granted, the initial point could also be the present (i.e. the end of the timeline). If participants provide their “when” answers accordingly, then the expectation would be that 4-item and 8-item conditions would not differ seeing that they were the same duration. Elaborating predictions of the central tendency for the nested effects reported here would require one additional assumption: that the scalar properties of item durations are rescaled with event boundaries. Making this assumption would then impose a new regression-to-the-mean of the scalar values per event. Doing so would now predict that the 4-item conditions would show a central tendency to the mean of 4 item positions (here, 8 s), whereas the 8-items conditions should be twice as high (16 s). The prediction of the central tendency rescaled to event duration predicts a steeper central tendency for the longer event (8-items) than for the shorter one (4-items).

A central tendency effect complemented with the assumption of a boundary reset predicts that the distances for the first items within an event would be overestimated relative to the last items within an event, which would be underestimated (following Vierordt’s or the central tendency effect). The corollary is that items before a boundary will be more distant relative to the following items belonging to the subsequent event than early items of an event will be from the last items of the previous event. The slope of the temporal compression should differ as a function of the number of items by event, with a larger compression predicted for larger events: here, however we find the opposite (Fig. 6). In summary, a pure central tendency effect would predict a global compression towards the midpoint of the sequence of 24 images without additional variations in temporal distances within or between events. Our data reveal an expansion of temporal distance towards the end of an event. A compression of distances between events and a global compression with more extreme compression for the last events than the first events. The quantity of information within nested event structures determines the ‘when’ of memory.

Memory for when depends on the reconstruction of information to establish similar distances between temporal contexts. Temporal displacements vary based on the compression of information at the level at which performance is tested and not on the objective number of seconds. Future work will examine finer-grained testing of memory for ‘when’ at different levels of nested event structures based on the accumulation, compression, and reconstruction of information building towards a nested compression model of temporal memory (Figure 6, Nested compression).

## CRediT author statement

**Matthew Logie:** Conceptualisation, Methodology, Software, Validation, Formal analysis, Investigation, Resources, Data curation, Writing – Original Draft, Writing – Review & Editing, Visualisation.

**Camille Grasso**: Investigation, Resources, Writing – Review & Editing.

**Virginie van Wassenhove:** Conceptualisation, Methodology, Investigation, Formal analysis, Resources, Writing – Review & Editing, Visualisation, Supervision, Project administration, Funding acquisition.

## Supporting information

Virtual reality task (Eight-item)

## Acknowledgements

We thank the UNIACT team at NEUROSPIN for assistance with data collection. We thank the members of the Cognition and Brain Dynamics team for helpful discussions, especially Marianna Lamprou-Kokolaki and Anna Wagelmans. We also thank Aidan Horner for helpful discussions.

## General Disclosures

All authors declare no conflicts of interest. Artificial intelligence: No artificial intelligence assisted technologies were used in this research or the creation of this article. Ethics: This research is in agreement with the declaration of Helsinki (World Medical Association, 2013) and the local Ethics Committee on Human Research (CEA-CPP-100049).

Data available at the following link: https://osf.io/4zqhc/overview?view_only=c93ed0a0ba75489a87225f6e4d063fea

No aspects of the study were preregistered.

